# *In silico* assessment of primers for eDNA studies using PrimerTree and application to characterize the biodiversity surrounding the Cuyahoga River

**DOI:** 10.1101/027235

**Authors:** M.V. Cannon, J. Hester, A. Shalkhauser, E.R. Chan, K. Logue, S.T. Small, D. Serre

## Abstract

Analysis of environmental DNA (eDNA) enables the detection of species of interest from water and soil samples, typically using species-specific PCR. Here, we describe a method to characterize the biodiversity of a given environment by amplifying eDNA using primer pairs targeting a wide range of taxa and high-throughput sequencing for species identification. We tested this approach on 91 water samples of 40 mL collected along the Cuyahoga River (Ohio, USA). We amplified eDNA using 12 primer pairs targeting mammals, fish, amphibians, birds, bryophytes, arthropods, copepods, plants and several microorganism taxa and sequenced all PCR products simultaneously by high-throughput sequencing. Overall, we identified DNA sequences from 15 species of fish, 17 species of mammals, 8 species of birds, 15 species of arthropods, one turtle and one salamander. Interestingly, in addition to aquatic and semiaquatic animals, we identified DNA from terrestrial species that live near the Cuyahoga River. We also identified DNA from one Asian carp species invasive to the Great Lakes but that had not been previously reported in the Cuyahoga River. Our study shows that analysis of eDNA extracted from small water samples using wide-range PCR amplification combined with high-throughput sequencing can provide a broad perspective on biological diversity.

## Introduction

Environmental samples, such as river and pond water or soil, contain a complex mixture of DNA molecules originating from intact microorganisms, feces, mucous, gametes, shed tissues or decaying parts from organisms living in or near the sampling site^1^. DNA extracted from these samples (often referred to as environmental DNA or eDNA) can be characterized globally by shotgun sequencing all DNA present within a sample. This approach (referred to as metagenomics) provides a wealth of information, not only about the identity of the species present in the environment, but also about the gene content of the sequenced organisms which can highlight interesting biological features^2, 3^. However, this approach requires extensive sequencing to capture the biological complexity of each sample and is therefore expensive to analyze many samples. Alternatively, DNA extracted from environmental samples can be analyzed by PCR targeting a carefully selected locus. This approach has been widely applied to ecological studies^4-6^, conservation^7, 8^ or to identify the presence of invasive species^9^. This approach is particularly cost-efficient since all reads generated are informative with regard to species identification, enabling detection of even rare organisms. Such eDNA studies typically focus on the analysis of a single macroorganism species and therefore rely on species-specific amplification of DNA (i.e., using primers that only amplify the species of interest). By contrast, characterization of microbial communities from environmental samples^10^ are often conducted using “universal” primers amplifying bacterial 16S ribosomal RNA genes (rRNA) that yield, after DNA sequencing, enough sequence information to identify the species carrying each DNA molecule.

Here, we use next-generation sequencing to characterize PCR products amplified from each eDNA sample using taxon-specific primers to obtain a quick and cost-efficient assessment of the macroorganism diversity. First, we describe PrimerTree, a novel R package that enables *in silico* evaluation of the specificity and information content of “universal” primer pairs. Next, we describe the application of this approach to the analysis of eDNA extracted from 91 samples of 40 mL of surface water collected along the Cuyahoga River (Ohio, USA). We amplified each sample using 12 primer pairs targeting mammals^11^, fish^12^, amphibians^12^, birds^13^, bryophytes^14^, arthropods^15^, copepods and plants^16^ (as well as several microorganism taxa^13, 17, 18^) and, after indexing, sequenced simultaneously the PCR products by massively parallel sequencing. Our analyses show that this methodology can provide a broad perspective on the aquatic and terrestrial biodiversity at the sampled sites in a simple, rapid and cost-effective manner.

## Materials and Methods

### In-silico evaluation of universal primer pairs

To evaluate the amplification breadth and informativity of “universal” primers we developed PrimerTree, an R package that performs the following functions for each primer pair provided by the user:

1. *In silico* PCR against a selected NCBI database
2. Retrieval of DNA sequences predicted to be amplified
3. Taxonomic identification of these sequences
4. Multiple DNA sequence alignment
5. Reconstruction of a phylogenetic tree
6. Visualization of the tree with taxonomic annotation

PrimerTree utilizes the *in silico* primer search implemented in Primer-BLAST^19^ by directly querying the NCBI Primer-BLAST search page. This allows access to all options available on the NCBI website. By default, PrimerTree searches the NCBI (nt) nucleotide database but alternative NCBI databases, such as only assembled genomes, or Refseq mRNA, can be queried. Note that when the proposed primers are degenerate, PrimerTree automatically tests up to 25 possible combinations (by default, with more possible) of primer sequences in Primer-BLAST and merges the results. The primer alignment results are then processed using the NCBI E-utilities (http://www.ncbi.nlm.nih.gov/books/NBK25500/) to i) retrieve DNA sequences located between the primers (i.e., the “amplified” sequences) and ii) obtain taxonomic information related to each DNA sequence using the NCBI taxonomy database (http://www.ncbi.nlm.nih.gov/books/NBK21100/). PrimerTree next aligns all “amplified” DNA sequences using Clustal Omega^20^ with a user configurable substitution matrix and reconstructs a Neighbor-Joining tree using the ape package^21^. Finally, PrimerTree displays the resulting phylogenetic tree using the ggplot2 package, labeling each taxon in a different color and adding the names of the main taxa using the directlabels package (http://CRAN.R-proiect.org/package=directlabels)^22^.

PrimerTree usually runs in less than five minutes, but the runtime varies greatly depending on the primer specificity (i.e., how many DNA sequences are “amplified”), the search parameters chosen, the current load on the NCBI servers and internet connection. In particular, each degenerate position in a primer will result in up to four times as many primers to be tested, which can considerably increase the runtime. To limit maximum runtime in this situation, PrimerTree randomly samples only a portion of the total primer permutations (25 by default). Changing the number of sampled permutations or including all variants is possible. PrimerTree uses the plyr package (https://cran.r-proiect.org/web/packages/plvr/index.html) extensively and has full support for any of the parallel backends compatible with the foreach package (https://cran.rproiect.org/web/packages/foreach/index.html). In particular, parallel retrieval of the primer sequences from NCBI speeds up the total runtime considerably. Note that parallel queries to Primer-BLAST are queued by NCBI’s servers and are only processed once there is free compute time.

### Sampling and DNA extraction

We collected water samples from the upper (n=39), middle (n=16) and lower (n=24) Cuyahoga River (**Figure 1** and **Supplemental Table 1** for details). These sections correspond to, respectively, an area with lower population density; a section with more human presence, dams and water treatment plants; and a heavily industrialized area. Each sample consisted of ∼50 mL of surface water collected roughly one meter from the bank of the river in a sterile 50ml conical tube. In addition, we also collected additional samples from water sources entering the river (n=12, **Supplemental Table 1**). The collections were performed during two sampling times separated by a major rain episode that dramatically increased the river discharge (**Supplemental Figure 1**). Samples were stored at ambient temperature until return to lab, where they were centrifuged and frozen at -20°C.

**Figure 1.**
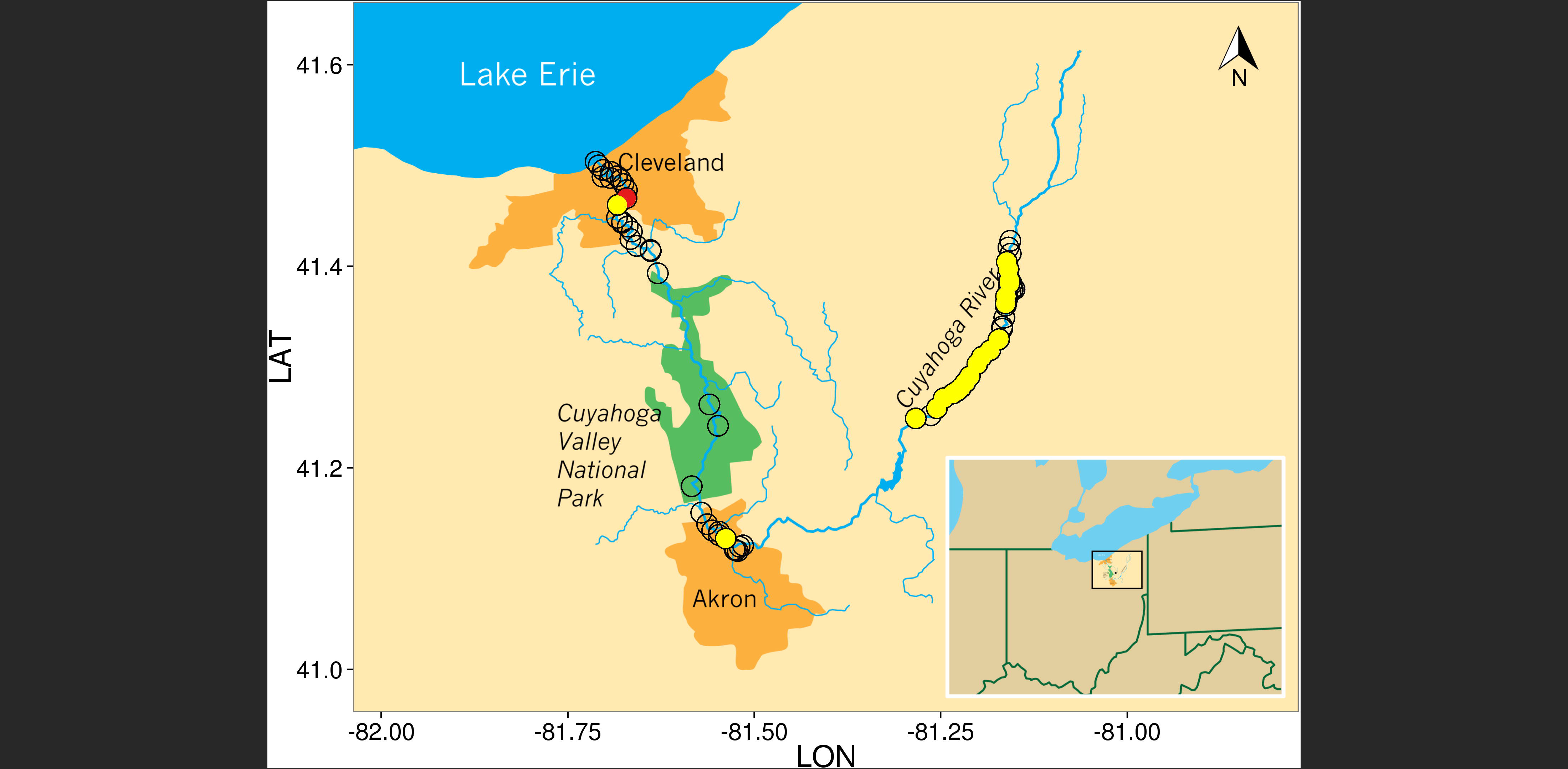
Geographic locations of water samples positive for Silver Redhorse and the Asian Carp eDNA using the 16S mammal primers. Each circle shows the location of a sampled site. The red circle indicates the location of the sample positive for Asian carp (*Ctenopharyngodon idella* or *Mylopharyngodon piceus).* Yellow circles are samples positive for Silver Redhorse (*Moxostoma cervinum* or *Moxostoma anisurum).* Map image was prepared by the Cleveland Clinic Center for Medical Art and Photography using Adobe Illustrator CS6 and points were overlaid using ggplot2.

We isolated DNA from each water sample using the following procedure adapted from previous studies^23, 24^. We first mixed each water sample by inversion and transferred 40 mL to a new tube for centrifugation at 8,000 x *g* for 30 minutes at 4°C. We discarded the supernatant and resuspended the pellet in 1 mL of ATL lysis buffer (DNeasy kit, Qiagen) supplemented with 0.47% Triton-X (Ricca Chemical Company), 7.88 mg of lysozyme (Fisher Scientific) and 19.2 units of lysostaphin (Sigma Aldrich). We then incubated the samples at 37°C for 1 hour while shaking them. We digested further by incubating 350 pi of each sample with 50 μl of proteinase K and 350 μl buffer AL (Qiagen) for 60 minutes at 56°C. Finally, we extracted DNA using Qiagen DNeasy columns according to the manufacturer’s instructions. We included two extraction controls and processed them identically and at the same time as the rest of the samples to monitor cross- or laboratory contamination. In addition, all experiments were performed in a laboratory where no eDNA or vertebrate DNA (aside from human and mouse) had been previously extracted or amplified.

### DNA amplification and sequencing

We selected primers to amplify different taxa from the literature and evaluated *in silico* their specificity and information content using PrimerTree. We required each primer set to amplify a region small enough for MiSeq reads to overlap to enable correction of sequencing errors. We amplified DNA extracted from each sample (with two extraction controls and one PCR negative control) using the following conditions: initial denaturation of 95°C for 15 minutes followed by 50 cycles of 95°C for 30 sec., 55°C for 30 sec. 72°C for 30 sec in 1X Quantitect mastermix (Qiagen) with 0.4 μM of each primer. Overall, we performed, on each sample, 12 independent DNA amplifications targeting Archaea (16S rRNA)^17^, Mammals (16S rRNA)^11^, Algae (23S rRNA)^25^, Amphibians (mt-Cytb)^12^, Birds (12S rRNA)^13^, Fish (mt-Cytb)^12^, Bryophytes (trnL)^13^, Arthropods (mt-Co1)^14^, Copepods (28s rRNA)^15^, Diatoms (18S rRNA)^18^, Fungi (ITS)^13^ and vascular plants (trnL)^16^ (**Supplemental Table 2**). Each primer included a 5’ tail for barcoding and Illumina sequencing (see below). We then pooled all 12 amplification products obtained from each water sample. We added Illumina adapter sequences and labeled each sample with an individual six nucleotide index (with each index distinct from all other index by at least two nucleotides) using primers targeting the 5’ oligonucleotide tail in 10 cycles of PCR (initial denaturation of 94°C for 3 minutes followed by 10 cycles of 94°C for 45 sec., 56°C for 45 sec. 72°C for 45 sec in 1X buffer, 1.25U GoTaq (Promega), 2mM MgCl_2_ and 2¼M of each primer). We then pooled all indexed samples together and we sequenced the resulting library on an Illumina MiSeq to generate 10,507,986 paired-end reads of 250 bp.

### DNA sequence analysis pipeline

We used custom PERL scripts to retrieve the index information and identify and trim the amplification primer sequences. We discarded any DNA sequence shorter than 50 bp, with the exception of sequences amplified with plant and bryophyte primers for which short amplification products were expected (see **Supplemental Table 1**). We also discarded any trimmed read pair for which the difference in sequence length was greater than 5 bp between the two paired-end reads to eliminate any reads where primers were not found in both reads. We then merged the paired reads into a single consensus DNA sequence using PANDAseq with default parameters^26^. We used Mothur^27^ to cluster unique DNA sequences and counted how many reads carried each unique DNA sequence.

While all raw sequences are freely available online (accession number SRP058316), we only describe here the analyses of macroorganism DNA sequences for sake of simplicity (microorganism sequences can be easily analyzed using standard packages such as those implemented in QIIME^28^). For macroorganisms such as mammals, many species have been sequenced for the locus of interest and if not, a closely related species is likely present in the NCBI database (but see also below). Therefore, to analyze DNA sequences from macroorganisms – mammals, amphibians, birds, bryophytes, arthropods, copepods and plants – we used BLAST^29^ to directly identify the closest DNA sequences in the NCBI database and the likely species of origin. Briefly, we removed from our analyses any DNA sequence observed in less than 10 reads total (summing across all samples), as these likely represent sequencing errors. We then compared each remaining DNA sequence to all sequences deposited in the NCBI nt database using Blastn (excluding uncultured samples) and only considered matches with greater than 90% identity over the entire sequence length. We then retrieved taxonomic data of all best match(es) for each sequence from NCBI. If multiple species matched a single sequence, all species names were assigned to the sequence. We conducted further analyses at the species level for all taxa, using a minimum read count per sample of 10 to determine absence/presence.

## Results

### In silico assessment of universal primer pairs using PrimerTree

PCR primers are usually designed to amplify one locus in a specific organism. Even “universal” primers, designed to amplify many species within the same taxonomic group, are typically used to only amplify DNA extracted from a single macroorganism (of this taxon). For studies amplifying eDNA or DNA from unknown taxa, we require that the amplification works on all members of a given taxon while avoiding off-target amplification (that could reduce the number of sequences from the desired taxon). In addition, the amplified region should contain enough sequence information to identify the species carrying the DNA sequence.

PrimerTree enables a rapid and visual assessment of these parameters for any primer pair by displaying the results of *in silico* PCR as a taxonomically-annotated phylogenetic tree. For example, **Figure 2** shows a subset of the PrimerTree results for a primer pair targeting the mammalian mitochondrial 16S ribosomal RNA genes (**Figure 2A**) and primers designed to amplify the chloroplast trnL gene of non-vascular plants (**Figure 2B**). The tree display enables rapid evaluation of the specificity of the primer pairs (e.g., off-target amplification of amphibians and ray-finned fishes on **Figure 2A**). In addition, the information content can be easily assessed by the length of the branches leading to different sequences (scaled in number of nucleotide differences). For example, PrimerTree reveals much longer branch lengths on **Figure 2A** than in **Figure 2B** suggesting a better discriminating power for the mammalian sequences than the bryophyte sequences (see also below). By default, PrimerTree displays phylogenetic trees annotated at all taxonomic levels enabling the user to determine the level of specificity of each primer set (**Supplemental Figure 2**).

**Figure 2.**
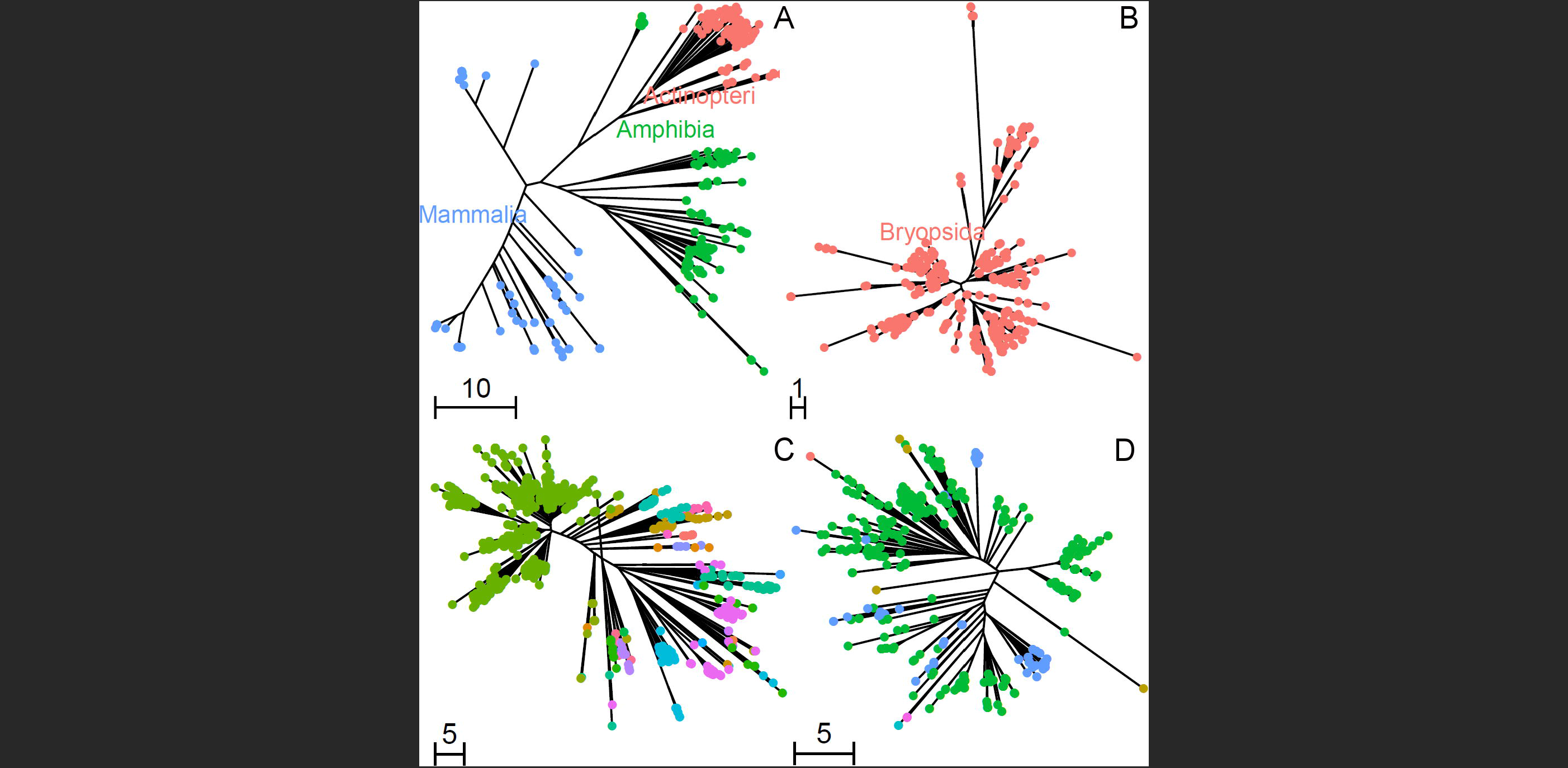
Example of PrimerTree results. The figure shows phylogenetic trees annotated at the class level for (**A**) the mammalian 16S rRNA and (**B**) bryophyte trnL primer pairs. The complete PrimerTree results for all 12 primers used in this study are presented in Supplemental Figure 2. Panels **C** and **D** are PrimerTree results from mammal 16S rRNA and fish mt-Cytb primers, respectively, demonstrating a greater breadth of amplifiable sequences by the mammal 16S rRNA primers. 1,000 BLAST hits within *Actinopterygii* are shown and tips are colored by taxonomic order. Fish mt-Cytb PrimerTree results include only 6 orders, while mammal 16S rRNA primers can amplify 27. The order *Cypriniformes* (which includes carp) is green in both panels.

### High-throughput sequencing of DNA amplified from river samples

We analyzed DNA extracted from 40 mL of surface water collected in 91 sites along the Cuyahoga River (**Figure 1**). For each sample, we performed 12 PCR amplifications targeting mammals, amphibians, birds, fish, arthropods, copepods, bryophytes and vascular plants (as well as several microorganism taxa not analyzed here). We then individually indexed the PCR products of each sample and sequenced them on an lllumina MiSeq (**Figure 3**) to generate a total of 10,507,986 paired-end reads (**Table 1**). After stringent quality filtering we retained between 1,645,452 and 6,213 reads for the analysis of each taxon (**Table 1**).

**Figure 3.**
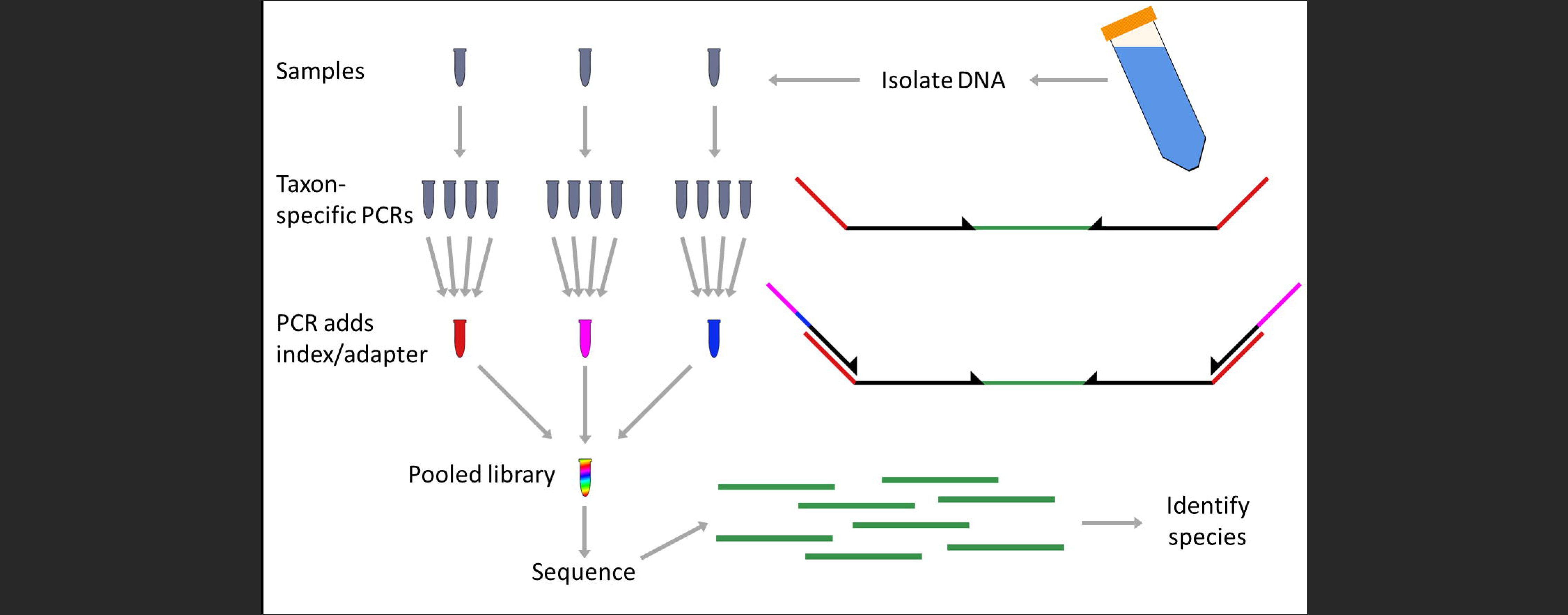
Experimental workflow. We first isolated DNA from 40 ml of river water. We then amplified each sample with 12 taxon-specific primer sets and pooled a portion of each PCR for each sample. Each primer had a 5’ tail to allow a second PCR which added Illumina adapter sequence and an individual index. We then pooled all barcoded samples and sequenced the library on a MiSeq. We used the sequence information to identify species of origin for DNA fragments isolated from the original samples.

**Table 1.** Sequencing and sequence analysis summary

| Taxon targeted | Locus | Reads generated | Reads after QC | Unique seq. | Unique seq. w/ >= 10 reads | Reads represented by unique seq. >= 10 | Species or OTUs identified | Seq. length (min-max) |
| --- | --- | --- | --- | --- | --- | --- | --- | --- |
| Mammals | mt16S rRNA | 1,384,149 | 1,352,524 | 13,238 | 786 | 1,328,072 (98.2%) | 65 | 95 bp (76 - 118) |
| Amphibian | mt-Cytb | 6,517 | 6,213 | 225 | 20 | 5,724 (92.1%) | 2 | 68 bp (68 - 68) |
| Birds | 12S rRNA | 79,549 | 64,144 | 873 | 136 | 62,433 (97.3%) | 123 | 53 bp (50 - 211) |
| Arthropods | COI | 299,714 | 42,987 | 5,043 | 238 | 34,855 (81.1%) | 25 | 162 bp (156 - 166) |
| Copepods | 28S rRNA | 502,972 | 79,486 | 13,252 | 368 | 59,753<br>(75.2%) | 5 | 207 bp (53<br>- 368) |
| Fish | mt-Cytb | 280,184 | 83,123 | 1,918 | 231 | 80,393<br>(96.7%) | 5 | 90 bp (76 -<br>192) |
| Bryophytes | TrnL | 253,053 | 225,780 | 1,728 | 262 | 223,153<br>(98.8%) | 32 | 52 bp (25 -<br>55) |
| Vascular<br>plants | TrnL | 1,954,381 | 1,610,089 | 33,584 | 3,468 | 1,554,086<br>(96.5%) | 236 | 48 bp (17 -<br>80) |

### Species identification and resolution

In contrast to microorganisms, where several hundred species are likely present in a given sample, we only expect to amplify DNA from a few different macro-organism species per sample. Therefore, even if the number of initial DNA molecules from a species is low, as long as the template can be amplified efficiently, many reads will be generated from these few DNA sequences. For example, if a sample had DNA from ten mammals and one species only accounted for 1% of the total DNA, its DNA sequence would be represented, on average, by 142 reads (mammal amplifications were represented by, on average, 14,211 reads per sample). We therefore considered that DNA sequences represented by less than 10 reads total across all samples were caused by sequencing errors and discarded them, removing between 1% and 25% of all reads generated, depending on the taxon considered (**Table 1**). We blasted the remaining DNA sequences to identify the closest DNA sequences in NCBI and to assign a species label to each DNA sequence.

In agreement with our *in silico* analyses, we observed large variations among primers in the specificity of the taxa identified (**Supplemental Table 3**). For some primers, the sequence information was insufficient to differentiate the organisms down to the species level: for example, each DNA sequence amplified from the trnL gene of vascular plants matched sequences from 34.89 different species on average (**Supplemental Table 4**). In fact, DNA sequences amplified from this primer pair matched a single taxon only when considering families or higher taxonomic levels. This contrasted with more informative DNA sequences such as the mammalian 16S rRNA for which each DNA sequence generated matched, on average, only 1.27 species (**Supplemental Table 3**). Note that the observed specificity of the bird primers differed from that expected from *in silico* analyses (12.39 species per DNA sequence generated compared to 1.57 expected based on sequences available in NCBI). This apparent low specificity in our data was caused by the presence of many DNA sequences from thrushes (a family of passerine birds) for which many species with the exact same DNA sequences at the 12S rRNA gene have been sequenced. In addition to differences in their information content (that influences the ability to identify a given sequence), primer pairs also differed in the amplification specificity. For example, while initially designed to amplify mammals^11^, the mammalian 16S rRNA primers also amplified *Actinopterygii* (ray-finned fishes). On the other hand, the primers targeting cytochrome B of fish^12^ only amplified common carp (*Cyprinus carpio*) and creek chub (*Semotilus atromaculatus*) under our PCR conditions as predicted by our PrimerTree analysis (**Figure 2C,D** **and Supplemental Table 5**). The “mammalian” 16S rRNA primers successfully amplified 31 samples for common carp while 15 were amplified using the cytochrome B primers, indicating a higher sensitivity (under our PCR conditions). The samples positive for common carp by the cytochrome B primers were usually also positive by the 16S rRNA primers: of the 15 samples positive for common carp using cytochrome B primers, 11 were positive using 16S rRNA primers (**Supplemental Table 6**). The four samples that did not yield common carp DNA using rRNA primers (but were positive using cytochrome B) could perhaps be explained by very low amount of fish DNA molecules (and stochastic amplification) or by the presence of much more abundant amplifiable DNA (e.g. mammalian DNA) that might have entirely swamped this signal. Consequently, for all subsequent analyses we used fish sequences amplified by the mammalian 16S RNA primers rather than those amplified with the cytochrome B fish primers.

### Molecular assessment of the biodiversity of the Cuyahoga River

Overall, across 91 water samples collected along the Cuyahoga River, we identified 54 samples positive for fish DNA (representing 15 species), 77 samples positive for mammalian DNA (17 species excluding human), 12 samples positives for bird DNA (from at least eight species), 18 samples positive for arthropod DNA (15 species), 16 samples positive for copepod DNA (two species) while the “amphibian” primers amplified turtle and two-lined salamander DNAs in two samples (**Table 2**, **Supplemental Table 6**). In addition to many organisms living in the river (e.g., fish, aquatic insects) or semi-aquatic animals (beaver, mink, muskrat), we also amplified DNA from many terrestrial species that live near the banks of the Cuyahoga River such as raccoon, groundhog, squirrel or mouse. Similarly, we identified DNA from many birds that live on (swan, duck, sea gulls) as well as near the river (sparrow, wild turkey). The species identified often corresponded with the local environment where the samples were collected: for example, beaver DNA was amplified from samples collected in protected forested areas, gull DNA near Lake Erie. In this regard, it is interesting to note that fish DNA showed significant differences in their geographical distribution. DNA from fish of the *Moxostoma* genus (probably Silver Redhorse) was commonly detected in the Upper Cuyahoga River but rare elsewhere (p=0.02, **Figure 1**). The central stoneroller was detected only in the middle and lower Cuyahoga (p=3.1x10^−3^). On the other hand, common carp were found throughout the entire Cuyahoga River (p=0.23). Surprisingly, we also identified DNA from one invasive Asian Carp species in the Cuyahoga River near Lake Erie (**Figure 1**). Note that the extraction and PCR controls almost exclusively yielded human DNA sequences (>99 % of the reads) with one extraction control also displaying pig DNA in 0.6 % of the reads. It is important to note here that, since the products of all PCRs were pooled and sequenced, regardless of the presence of detectable amplified products on an agarose gel, the sensitivity of this approach is magnified compared to standard molecular approaches and laboratory contamination with human DNA difficult to avoid. However, these controls indicated that DNA sequences (aside from human sequences) retrieved from the water samples must be genuine and that cross-contamination in the laboratory was minimal.

**Table 2.**
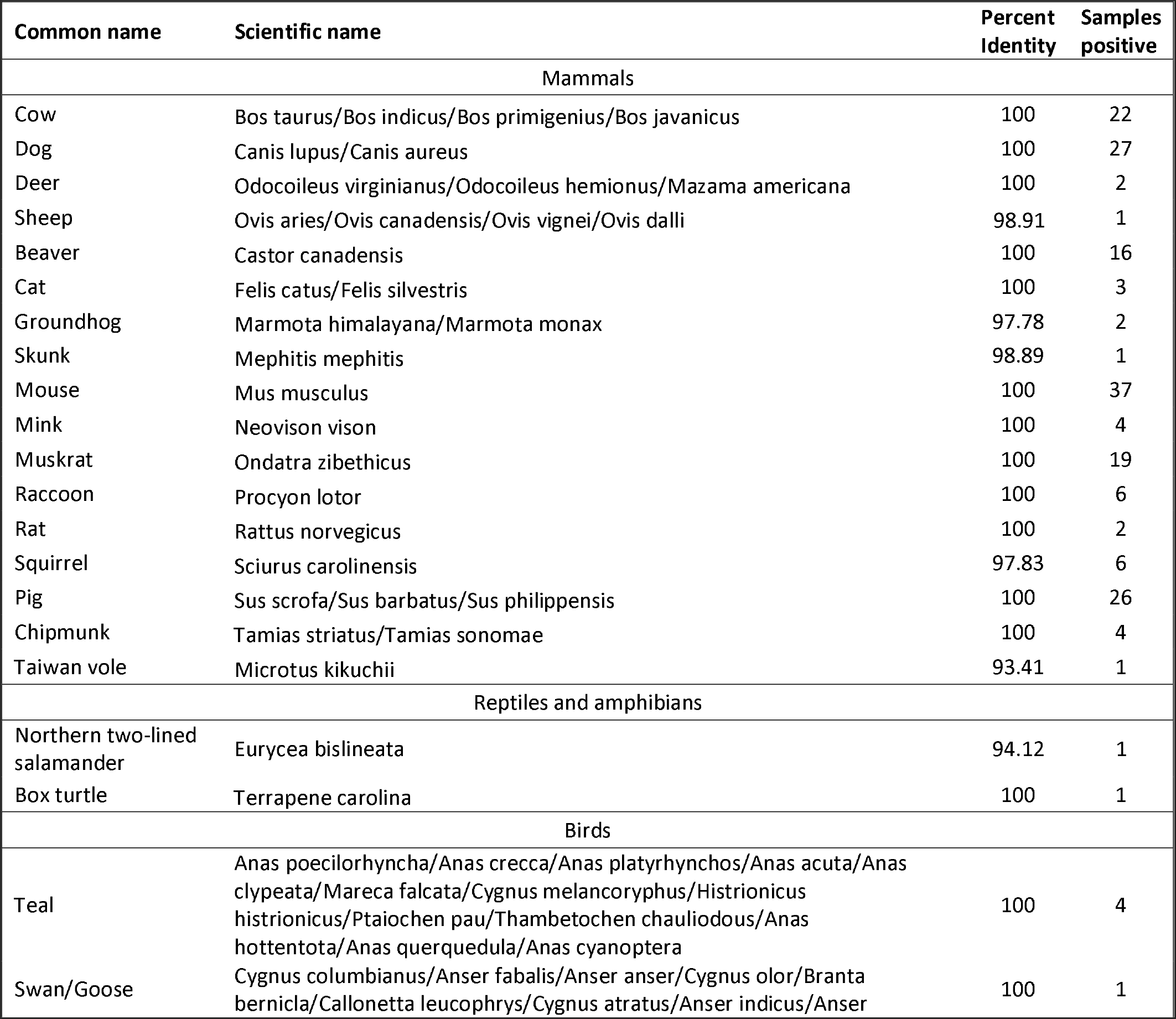
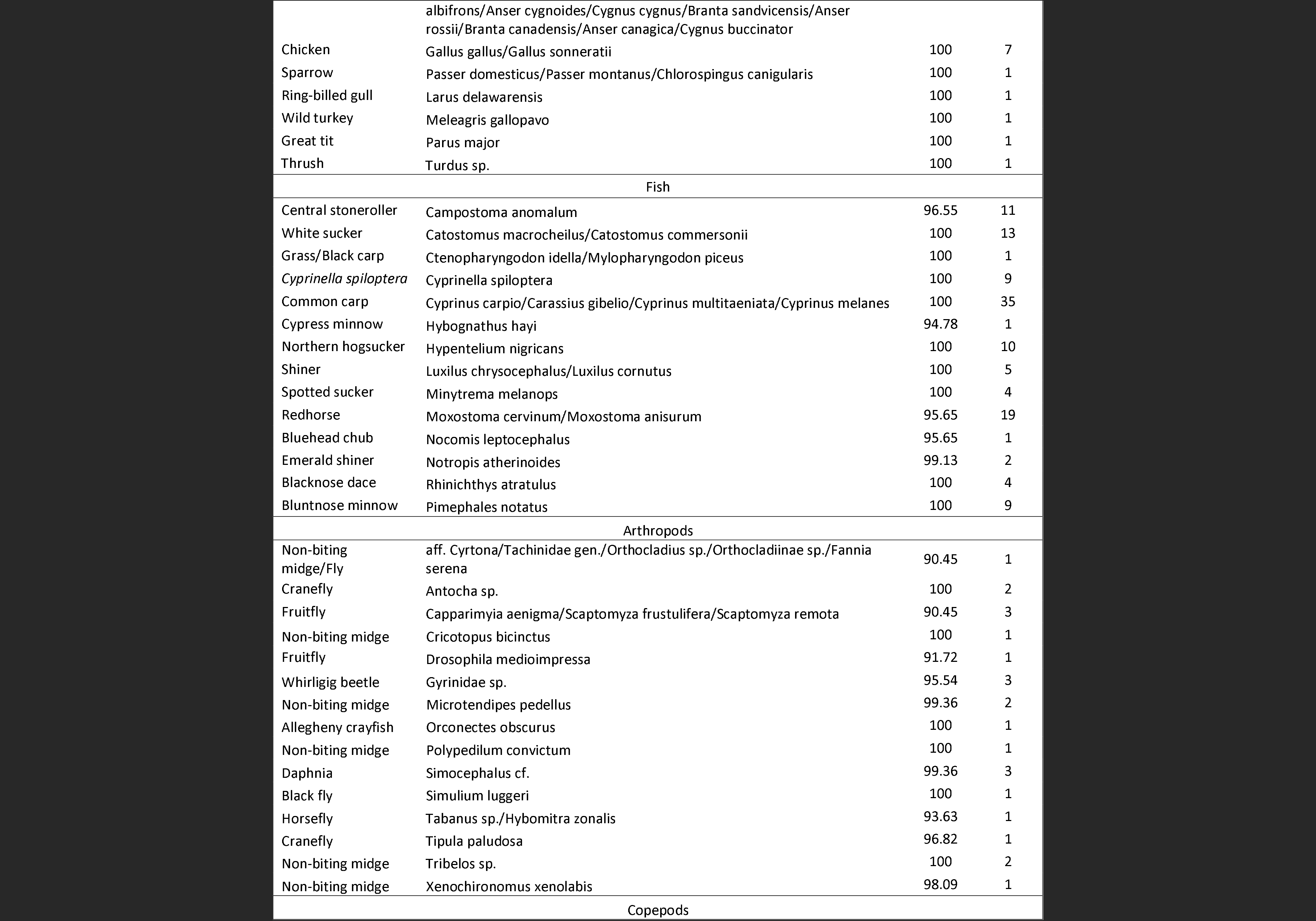
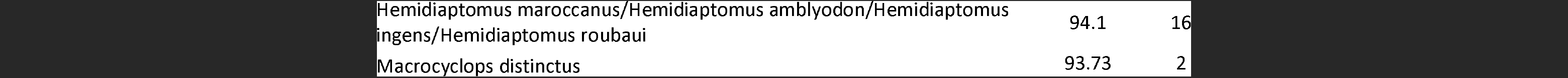
Macro-organisms identified in the Cuyahoga River.

### Temporal variations in diversity

We performed the collection of water samples at two time points separated by an episode of heavy rain falls that dramatically altered the water level and flow of the Cuyahoga River: the discharge at Hiram Rapids, in the Upper Cuyahoga River, increased from 2.46 cubic meters per second (close to the median daily statistics) at the time of the first sampling to 20.87 cubic meters per second and was still as high as high as 5.95 cubic meters per second at the time of the second sampling 12 days later (**Supplemental Figure 1**). For most macro-organisms, we did not detect any statistical difference between the samples collected at the two time points. One notable exception concerns fish DNA: we observed significantly more positive samples before rain than after (p=0.05, Fisher’s exact test).

## Discussion

Since the first reports that DNA could be retrieved from environmental samples^30^, studies of environmental DNA have broadened in scope from studies of bacterial communities (e.g., ^2, 31^), to identification and monitoring of a given species (e.g., ^5, 9, 12, 32^) and the characterization of microorganism populations (e.g., ^33, 34^) and recently to the identification of macroinvertebrates or vertebrates (e.g.,^35-38^). However, several factors have limited a broader implementation of these approaches for ecological studies. These limitations include the lack of tools enabling a wide range of species to be studied simultaneously, the amount of starting material (often liters of water for aquatic environments) and the high costs of such analyses. Additionally, the careful evaluation of potential primer pairs prior to laboratory work is critical as most published primers have only been tested on DNA directly extracted from the target organism and, while efficient, might not be specific or could even better amplify other organisms.

### PrimerTree provides a robust assessment of primer pairs for eDNA studies

We present here a simple R package that enables rapid screening of primers suitable for differentiating species from a chosen taxon. PrimerTree allows evaluating the specificity of a given primer pair (to avoid off-target amplification) and whether the amplified DNA sequences would provide enough information to identify the organisms carrying the DNA sequences. Since the amplified regions are typically short (100-300 bp), the resulting phylogenetic trees (**Figure 2** and **Supplemental Figure 2**) do not necessarily reflect the true species relationships, but they enable an easy and rapid assessment of the primer suitability. First, the automatic taxonomic annotation of the branches allows the user to quickly see which taxa present in the BLAST database are amplifiable by the primers. This enables identifying that undesirable (off-target) taxa may be amplified or that the primers might not amplify specific genera within the taxon of interest. Second, the branch lengths show how many nucleotides separate DNA sequences from different species. Short branch lengths might results in a low resolution in taxon identification (or elevated risk of false-positives caused by sequencing or PCR errors) while long branches will lead to high confidence species identification (e.g., **Figure 2A** vs. **2B**).

An attractive feature of PrimerTree is its ease of use. As an R package, it is easily installed or updated and only requires that clustal be installed in the user’s path. The entire process of getting BLAST hits for a primer pair is done through R using a single command and plots are generated using a second command. The summary function provides many useful statistics including amplified DNA sequence lengths, number of taxa amplified and average pairwise differences within each taxonomic level. Additionally, information on the BLAST results, sequences obtained, taxonomy of amplifiable sequences, neighbor joining distance matrix and phylogenetic tree are preserved within the PrimerTree object if the user wants to further analyze or further summarize the results. Finally, PrimerTree runs fast: for a primer pair with no degenerate bases, PrimerTree results are usually obtained in less than five minutes.

### Limitations of PrimerTree and comparison with other programs

One limitation of PrimerTree is that, by default, it only retrieves up to 500 random amplifiable sequences from the NCBI database and this random subset might not accurately represent the specificity of a given primer pair for a particular environment. For example, the bird primers used in our study display a high species-specificity for most birds but were not able to differentiate among thrushes. Note however that this issue can be easily circumvented by selecting specific targets in PrimerTree if one is interested in a particular taxon (e.g., one could run PrimerTree for querying only *Turdidae* sequences in NCBI to evaluate if a given primer pair is informative within this taxon). Additionally, the problem of returning a random subset of all sequences can be minimized by requesting more amplifiable sequences using the num_aligns argument in the command. Another important limitation is that PrimerTree does not highlight taxonomic groups that are not retrieved. If a particular taxonomic group is not amplifiable by a primer pair, the user must recognize the absence of that taxon. For example, if one primer pair amplifies all mammals except monotremes, the user must note the absence of the order *Monotremata* from the resulting plot or taxonomic information in the PrimerTree object. Along these same lines, the absence of a species or larger taxonomic group from PrimerTree results may not mean that the assayed primers cannot amplify those taxa. As PrimerTree uses the BLAST nr/nt database as a reference, any species without DNA sequence for the targeted locus in the database cannot be retrieved.

EcoPCR^39^ is another program than can evaluate primer specificity by utilizing a user-provided database to identify amplifiable sequences and provides a summary of amplifiable sequences and their taxonomic information. The ecoPCR program retrieves all amplifiable sequences in a custom database, compared to only a subset of the BLAST nr/nt database for PrimerTree. However, one advantage of PrimerTree is that it always queries the most current available version of the entire BLAST nr/nt database, while the user would need to update the database provided to ecoPCR and re-generate the ecoPCR database files to include new sequences in the analysis. Additionally PrimerTree benefits from the graphical output, which is automatically generated using the plot function in R, to summarize the results in a user-friendly manner.

### Characterization of eDNA extracted from the Cuyahoga River

We analyzed eDNA extracted from 91 samples of water collected along the Cuyahoga River. We showed that, despite a small amount of starting material, we were able to retrieve eDNA from a wide range of organisms, including many vertebrate species: on average, each water sample contained eDNA from 3.2 vertebrates (fish, mammal or bird), 0.3 arthropods and numerous plants. The most exciting feature of our findings is the amplification of DNA from many terrestrial organisms from the river samples. We amplified DNA sequences from deer, squirrel, raccoon, groundhog, vole, mink and skunk in addition to semi-aquatic species such as beaver and muskrat. We also identified DNA sequences for agricultural species (cow and swine) and companion animals (dog and cat), consistent with the findings of a study recently made available through bioRxiv (http://dx.doi.org/10.1101/020800). DNA from these species in the river samples likely originated from fecal matter washing into the river, deceased animals in or around the river or animals drinking or wading in the river. Similarly, we detected many avian sequences, including species that do not live in the river (e.g., chickens, turkeys, blackbirds and sparrows).

Several species were unevenly distributed along the Cuyahoga, such as the Silver Redhorse that was mostly detected in the Upper Cuyahoga, which demonstrates the potential of our approach to identify variations in species distribution (**Figure 1**). Among the 48 water samples positive for fish DNA, we also identified one sample containing DNA from one Asian carp (either *Ctenopharyngodon idella* or *Mylopharyngodon piceus)*, invasive species of the Great Lakes that were not known to be present in the Cuyahoga River (**Figure 1**).

### Biological and technical factors influencing species detection and the rate of false negatives

Our two samplings of the Cuyahoga River were separated by a major rain event. Since we sampled the upper portion of the Cuyahoga both before and after the rain, we compared species detection rates to determine if the influx of water into the river would stir up sediment and wash genetic material into the river to increase species detection rates or dilute the river water and decrease species detection rates. We found that the number of samples positive for mammals, birds, arthropods and copepods did not change after the rain, whereas the number of samples positive for fish species decreased after the rain.

This may suggest that, for fish, the rainwater dilutes the genetic material of fish, but for other taxa this dilution effect might be balanced by the influx of genetic material in water washing into the river. Replication of these results using other sample sets is necessary to confirm that this is a general phenomenon.

Certain taxa were identified less frequently than we would have expected (e.g., amphibians, arthropods and copepods, **Table 1**). This could be due to a number of factors. Our DNA extraction protocol (relying on the DNeasy extraction kit after centrifugation) may not have been optimal to retrieve DNA from such organisms as the choice of DNA isolation protocol from water samples can strongly influence the proportion of DNA isolated from different taxa^40^. In particular, it is important to note that our isolation method was unlikely to capture free DNA in the river water sample, but rather only DNA within, or adhered to, particulate matter which could lead to disproportional representation of the eDNA (see e.g., ^24, 41^. It is also possible that the PCR conditions we used were inadequate to efficiently amplify some of the targeted templates^42, 43^. Alternatively, the amount of genetic material present in the river may be lower for these taxa than for others such as fish or mammals (possibly due to differences in the size of the dead animals or the amount of feces or shed tissues). Additional studies, including experimental validation of the primers for these taxa, will be necessary to differentiate these possibilities.

While detection of eDNA is by nature stochastic, the small sample size used in our study likely increases this randomness and increases the chance of false negatives. By optimizing PCR conditions and increasing the number of cycles, one can limit the chance of having a DNA template present in a given sample failing to amplify. Nonetheless, a small sample of water may not entirely capture the diversity present in the environment, especially for species represented by few DNA molecules in the environment. While the amplification of a species’ DNA sequence is a clear evidence of the presence of this species (assuming a low level of cross-contamination), the failure to detect an organism is not a proof of its absence in the environment as the false negative rate is likely to be important. For example, while 20 out of 38 samples in the upper portion contained common carp (*Cyprinus carpio*) DNA, we would not be able to rule out the presence of common carp at the other sampled locations in this part of the river. The use of universal primer pairs also contributes to the likely high false negative rates: since our assay is based on primers that amplify multiple taxa rather than being species-specific, it is possible that DNA sequences from one species completely overwhelmed the signal originating from rarer eDNA templates present in the same sample. This effect could be magnified if there are differences in amplification efficiencies^42, 43^. For example, it is possible that some samples that did not yield common carp sequences actually contained common carp eDNA but that these molecules remained undetected due to the abundance of other templates (e.g., Silver Redhorse or human) in these samples. The presence of multiple amplification targets is also one reason why this assay is non-quantitative: the number of reads obtained for one species in a given sample is not only determined by how much DNA from that species is present in the sample, but also by how many other amplifiable species are present. Therefore, one cannot directly compare species read counts between samples to determine relative abundance^36^. In these regards, it is important to emphasize that the present approach does not replace classical eDNA studies targeting a single species but is designed for enabling broader ecological survey (that may guide further in-depth investigations).

### A fast, high-throughput and cost-efficient method to characterize biodiversity from environmental samples

We described here a customizable approach that builds on existing methods^42-45^ to enable simultaneous analyses of eDNA extracted from many samples for a wide range of taxa.

We showed that this methodology can be successfully applied to analyze small water samples. This aspect, while it has its limitations (see above), is essential for any study for which sampling and storing of large volumes is logistically challenging or impossible. In particular, small volume collection enables sampling in sites that can only be reached by hiking or paddling and will therefore make this approach suitable for many ecological studies. Additionally, sample processing (such as filtration) in the field can be slow and requires specialized equipment, particularly for large sample volumes. Here, we chose instead to centrifuge the samples upon return to the laboratory. This approach isolates the same portion of the sample as filtration (particulate matter), is rapid and reduces potential sample contamination during filtration. Additionally, by minimizing in-field handling, more samples can be collected across an aquatic system in a given timeframe, minimizing temporal sampling bias. For instance, we were able to sample roughly half the length of the Cuyahoga River in a single day with a single sampling team using kayaks. The speed of sampling is also an important parameter as it enables studies investigating the consequences of specific events such as rainfall, ecological disasters (e.g., oil, chemical or pollution spills) or even transient ecological events such as fish spawning. However, depending on the specific goals of a study and the resources available, larger sample volumes or alternate DNA isolation methods might be preferable.

Finally, this approach is cost-efficient: in our study of the Cuyahoga River, we characterized eDNA amplified from mammals, fish, birds, amphibians, arthropods, vascular plants, bryophytes and many microorganisms in 91 water samples for a total cost of approximately US $2000 (including DNA extraction, PCR reagents and sequencing costs, see **Supplemental Table 7** for details). This high level of multiplexing (across samples and across taxa) decreases the burden of the next-generation sequencing price, but, thanks to the tremendous sequencing output, still provides enough reads to rigorously characterize the composition of each sample. The use of an inexpensive PCR to add the sequencing and barcoding adapters also dramatically reduces the cost compared to generation of typical next-generation sequencing libraries^18, 32, 36, 46^. Note that the cost and efficiency could potentially be further improved by multiplexing primer pairs in the PCR reaction^44^.

### Limitations and cautionary notes for future studies

One important limitation of our approach regards the interpretation of the results. First, because we performed the species identification of sequences using BLAST, it is affected by the content of the NCBI database and the reliability might vary for different taxa. For instance, our analyses revealed, in one sample, the presence of DNA sequences most similar to a Taiwanese vole, which is not present near the Cuyahoga River, but with only 93.4% identity. These sequences likely originate from a local species of vole (closely related to the Taiwanese vole) that has not been sequenced for the 16S rRNA gene. This illustrates that matches with low percent identity BLAST hits need to be cautiously interpreted. Using primer pairs targeting different loci (e.g., COI) could partially circumvent this limitation as one species may have been sequenced at one locus but not another. On the other hand, this example also shows that this method can be used to identify organisms even if they have not been previously sequenced for the locus of interest, as long as a closely related species is present in the database, but that rigorously identifying the actual species present will require further analyses. In addition, when considering best match for a given sequence, the identification should be considered in the context of the other sequences amplified from the sample. For instance, in our analyses we identified sequences that best matched unexpected fowl species (e.g., *Gallus lafayetii* and *Gallus sonneratii).* However, these DNA sequences were only observed in small numbers and in samples that had a much larger numbers of *Gallus gallus* sequences. The most likely explanation for these sequences is that these reads derived from PCR or sequencing errors from *Gallus gallus* templates: the chicken sequences were so abundant in these samples that even rare errors led to a number of reads sufficient to pass through our stringent filtering criteria. The same phenomenon generated a handful of sequences most similar to monkey or ape sequences in samples with many human reads.

One final issue is the risk of false positives generated by contamination: since PCR products are directly sequenced by massively parallel sequencing, even minute contamination can be detected and analyzed. It is therefore critical to implement stringent measures to prevent and detect contamination, similar to those used for ancient DNA studies (e.g., numerous negative controls, use of sterilized room and equipment, etc), to prevent cross-contamination and obtain reliable results.

### Conclusion

The study of environmental DNA using massively-parallel sequencing technologies enables characterization of species biodiversity in a simple, high-throughput and cost-effective manner. Our results reveal that a sampling protocol relying on small sample volume enables the preliminary evaluation of complex environments. The ability to characterize a very diverse range of taxa from a single sample in a high-throughput manner will allow future studies to expand the scope of the biodiversity studied and to explore complex ecological interactions among species. Additionally, the identification of eDNA of local terrestrial flora and fauna in river samples provides a simple way to assess the local diversity of environments adjacent to rivers or other water bodies. Overall, our findings illustrate the sensitivity and utility of broad surveys of eDNA by deep sequencing and shows how this approach can constitute an excellent foundation for ecological and environmental studies.

## Acknowledgments

This work was supported by Cleveland Clinic funds to DS.

## Author Contributions

MVC, STS and DS conceived the study. MVC, JH, AS, ERC, KL, STS and DS contributed to the sample collection. AS performed the DNA extraction and PCR amplification. MVC analyzed the data. JH implemented PrimerTree. MVC and DS drafted the manuscript. All authors have read and approved the final version.

## Data Accessibility

All sequencing data generated in this study are available through the SRA at NCBI (Accession #SRP058316 (data to be released upon publication)). The R package PrimerTree is freely available at http://github.com/jimhester/primerTree.

